# Monitoring terrestrial vertebrates with airborne DNA in the Luangwa Valley, Zambia

**DOI:** 10.64898/2026.03.11.711018

**Authors:** Daniel Gygax, Sabina Ramirez, Michael Riffel, Jan-Dieter Ludwigs, Geofrey Zulu, Tom Riffel, Fabian Roger, Lara Urban

## Abstract

Vertebrates play vital roles in maintaining ecosystem processes and services and serve as valuable indicators of environmental health, making them an important target for monitoring and conservation efforts. Within the environmental DNA (eDNA) toolbox, airborne environmental DNA has recently emerged as a novel approach for vertebrate monitoring. In this study, we evaluated on-site airborne eDNA for terrestrial vertebrate monitoring in the Luangwa Valley savanna in Zambia, which represents a major biodiversity stronghold of largely intact wilderness and with high levels of vertebrate diversity and endemism. Six air samplers were deployed over four days alongside camera traps for validation, and samples were processed using a mobile molecular laboratory. In total, 120 terrestrial vertebrate taxa were detected with airborne eDNA, including 16 of the 17 taxa recorded by camera traps, demonstrating high sensitivity. Notably, 72.5% of taxa were detected on the first day, and a single sampler recovered 61.7% of all taxa; the taxonomic richness incrementally increased with extended sampling efforts, but the magnitude of these increases declined progressively. The detected taxa spanned the four terrestrial vertebrate classes and encompassed a wide range of ecological traits. These results show that airborne eDNA can quickly recover a substantial and representative fraction of local vertebrate diversity within a short sampling window, while extended sampling can improve detection of less common taxa. Despite existing limitations, our findings support the use of airborne eDNA as an efficient and scalable complementary tool for community-level biodiversity assessments in terrestrial ecosystems such as Zambezian savannas.

## Introduction

Global biodiversity is declining, threatening ecosystem stability and the services nature provides (Ceballos et al., 2015; IPBES, 2019). Effective biodiversity monitoring is therefore essential to guide conservation actions and promote sustainable land management (Butchart et al., 2010; Schmeller et al., 2017; Jetz et al., 2019; Ruckelshaus et al., 2020). Vertebrates play vital roles in maintaining ecosystem processes and services and serve as valuable indicators of environmental health, making them an important target for monitoring and conservation efforts (Whelan et al., 2008; Ceballos et al., 2020; Gable et al., 2020; Yeakel et al., 2020; González-Varo et al., 2021).

Zambia is among the continent’s most biodiverse countries (Siachoono, 2018), with the Luangwa Valley in the Eastern Province, spanning over 40,000 km² of pristine wilderness, standing out as a key ecological region (Astle et al., 1969). The valley serves as a stronghold for some of Africa’s most iconic vertebrate species (Riggio et al., 2013; Anderson et al., 2015; Chabwela et al., 2017). It also supports several endemic and near-endemic species (Pocock, 1897; Cotterill, 2000; Coimbra et al., 2021) and is recognized as an important avian region, with around 450 bird species recorded (Gerkmann et al., 2008; Dowsett & Leonard, 2009). Despite its high ecological value and prominence for safari tourism, the Luangwa Valley faces growing anthropogenic pressure (Abel & Blaikie, 1986; Jachmann & Billiouw, 1997; Phiri et al., 2024). It is therefore important to develop adequate and evidence-based conservation management strategies that balance wildlife and local livelihoods, for which biodiversity monitoring is essential. (Bradshaw, 2004; Jadhav & Barua, 2012; Chidakel et al., 2021; Stephenson et al., 2022; Zyambo et al., 2024).

Traditional biodiversity monitoring techniques, such as transect surveys, camera trapping, or acoustic monitoring, have long been established as fundamental tools for assessing vertebrate species occurrence and abundance (Zwerts et al., 2021). However, these methods often face limitations related to detection bias, incomplete species identification and taxonomic coverage, and restricted geographic accessibility (Newey et al., 2015; Stephenson, 2020). Environmental DNA (eDNA) offers a toolbox of complementary methods that can help address these challenges (Kelly et al., 2014; Ruppert et al., 2019; Johnson et al., 2019). Among these, water eDNA is one of the most widely used and well-established sources of eDNA (Deiner et al., 2017; Ruppert et al., 2019). Natural ponds have been successfully used to monitor vertebrate communities across diverse ecosystems, including those in Africa (Harper et al., 2019; Schenekar et al., 2024). For example, a water eDNA study in the Luangwa Valley detected several terrestrial vertebrate species, which were complementary to the species detected by traditional camera trapping (Gygax et al., 2025).

Within the same toolbox, airborne eDNA has recently emerged as a novel approach for vertebrate monitoring (Clare et al., 2022; Johnson et al., 2023; Lynggaard et al., 2022, 2024; Polling et al., 2024; Cai et al., 2025; Craine et al., 2025). A recent study in the Netherlands showed that all vertebrate species detected by camera traps were also identified using airborne DNA, indicating high sensitivity of this method (Polling et al., 2024). Airborne eDNA also appears to provide a strong and localized signal, as suggested by studies on invertebrates (Roger et al., 2022), vertebrates in zoo environments (Lynggaard et al., 2022; Cai et al., 2025; Bodawatta et al., 2025), and single species detections (Kroos et al., 2026). This localization, likely due to dilution, rapid degradation, and limited particle transport, can be considered a positive feature for precise vertebrate monitoring due to an increase in spatial resolution in comparison to water eDNA (Roger et al., 2022). Airborne DNA can also offer scalability, with the opportunity to choose between high-capacity, long-term air sampling platforms and more portable, simple, and cost-effective samplers for short-term sampling, which allows for rapid deployment and easy transport (Tulloch et al., 2025).

Most airborne eDNA studies have used either Illumina (e.g., Lynggaard et al., 2022, 2024; Polling et al., 2024) or Oxford Nanopore Technologies’ (ONT) GridION sequencing devices (Craine et al., 2025). These sequencing devices have a restrictively high initial investment cost and are often restricted to specialized laboratories and sequencing facilities. In contrast, smaller and more affordable sequencing devices such as ONT’s MinION sequencer can facilitate fast on-site access to sequencing in low- and middle-income countries (LMICs) and remote areas, obviating the need for sample storage and transport (Pomerantz et al., 2018; Urban et al., 2023; Gygax et al., 2025).

In this study, we evaluated the potential of airborne eDNA metabarcoding for monitoring terrestrial vertebrates in Zambia using a fully portable laboratory setting. We assessed the extent of vertebrate diversity that can be detected through on-site airborne eDNA analysis in a subtropical savanna ecosystem. We then validated eDNA-based vertebrate detections by comparing results with camera trap records obtained in the immediate vicinity of the air sampling locations. We thus demonstrate the feasibility of implementing a complete airborne eDNA workflow, from sample collection to data analysis, under remote field conditions using a mobile laboratory.

## Materials and Methods

### Study area

Sampling was conducted around Mwende Lagoon (12.53318 S, 32.14350 E) in Luambe National Park (NP) within the Luangwa Valley in eastern Zambia. The lagoon is ∼260m long and ∼62m wide at its widest point and lies adjacent to a large dirt road (Figure 1). The vegetation surrounding the lagoon consists of Zambezian/Mopane savanna, one of several habitat types forming the ecological mosaic of Luambe National Park (Anderson *et al*., 2016). Because the lagoon holds water for most of the year, it functions as an essential dry-season refuge and water source for wildlife such as elephants, hippopotamuses, giraffes, antelopes, and a variety of bird species. Alongside semi-aquatic species such as crocodiles and frogs, its high faunal diversity makes it an ideal location for testing eDNA-based monitoring approaches.

**Figure 1.**
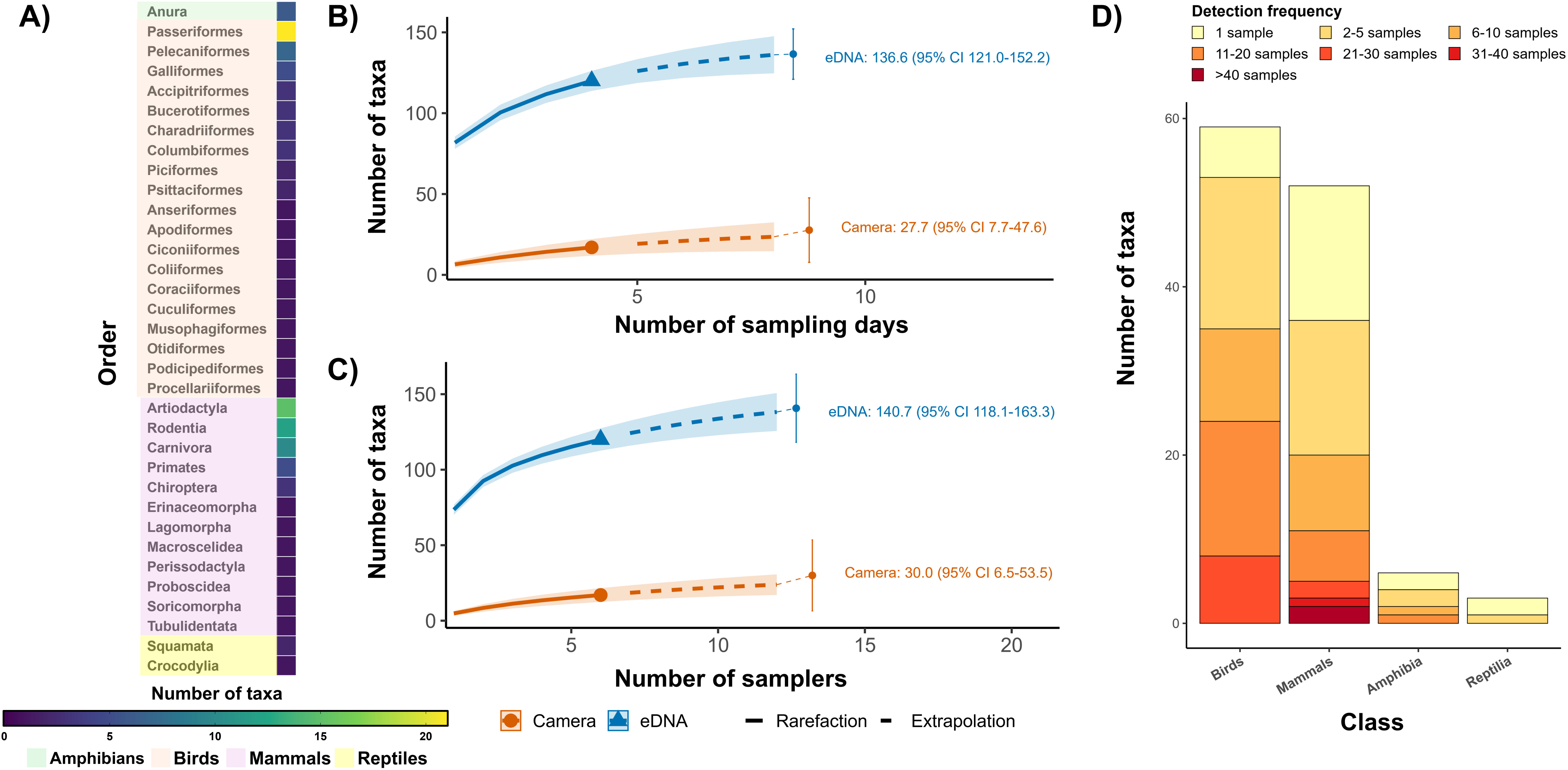
Location of the study area and sampling sites. A) Geographical location of Mwende Lagoon (Mw) around Luambe National Park, Zambia. Map created in QGIS using Esri Satellite imagery and Esri National Geographic basemap. Source: Esri, Maxar, Earthstar Geographics. B) Location of sampling sites (n=6) around Mwende Lagoon. C) Air Sampler 1 (see Methods). D) Panoramic view of Mwende Lagoon.

### Air sampling

Sampling of air took place between July 4^th^ and 9^th^, 2025. Six samplers were placed around the lagoon to provide coverage across most of its perimeter. Samplers were strapped to tree trunks at a height of 1.5m in locations that were readily accessible and free from dense vegetation.

Following every 24-hour sampling period, each sampler was disassembled, the filter was carefully removed with sterile tweezers, placed in a sterile Falcon tube, and immediately stored on ice until frozen in camp at −20 °C. The sampler was then thoroughly cleaned with 1% bleach before inserting a new filter. All procedures were conducted while wearing FFP2 masks and sterile gloves, which were regularly cleaned with bleach and exchanged between samplers.

Air sampling was conducted using compact, portable air eDNA samplers provided by DNAir AG (Zürich, Switzerland). The devices collect airborne eDNA particles onto sterile synthetic fiber filters by generating a continuous airflow of approximately 200 L min⁻¹ using an integrated Brushless Direct Current fan. Samplers were powered by commercially available 20,000 mAh power banks housed within an internal, rain-protected compartment, enabling autonomous operation for approximately 24 hours per deployment. At this flow rate and sampling duration, each sample represents a filtered air volume of approximately 288 m³. The sampler body (excluding battery) weighs <500 g, facilitating field deployment.

### Camera trapping

A camera trap (SECACAM Wild-Vision Full HD 5.0) was deployed at each sampling location to validate species detections. The cameras, equipped with passive infrared sensors, were set to photo mode and operated continuously throughout the air sampling period. Each camera was positioned to capture the most likely animal movement routes and to maximize the field of view, though not necessarily aimed at the air samplers. The photographs were manually reviewed by trained observers to identify all recorded species.

### DNA extraction

The synthetic fiber filters were placed inside a sterile 10 mL BD syringe barrel using sterile tweezers. Buffer ATL (3 mL) and Proteinase K (50 µL; 20 mg/mL) (QIAGEN, Hilden, Germany) were added directly onto the filter. Air was then expelled from the syringe barrel by advancing the plunger until the liquid fully filled the chamber, ensuring complete contact between the buffer and the filter. Afterwards, the syringe was sealed, and the contents were mixed by repeatedly moving the plunger and vortexing for 15 s. The syringes were then incubated upright with the cap facing down at 55 °C for 3 h. In the absence of a laboratory incubator or thermomixer, temperature was maintained using an aluminum kitchen pot filled with water and heated by a kitchen immersion circulator. After incubation, the ∼2ml lysate was expelled over a 2 mL low-bind tube. The lysate was centrifuged at 12 000 × g for 1 minute to pellet any particulate matter and clarify the supernatant and 600µl of lysate was transferred to a fresh 2 mL tube. After this, the Qiagen Blood and Tissue extraction protocol was followed with adjusted volumes (600 µl Buffer AL and 600 µl ethanol) until all the lysate was processed. DNA was eluted in 2 x 50 µl elution buffer except for samples from day 2 which were only eluted in 50 µl once due to a temporary centrifuge malfunction. DNA concentrations were measured with a Fluorometer (invitrogen™ Qubit™ 3.0) using the High Sensitivity (HS) Assay and purity ratios were measured with a Thermo Fischer™ NanoDrop One. After a PCR amplification attempt, samples were cleaned and concentrated using the Zymo™ DNA Clean & Concentrator-5 kit, with a final elution volume of 35 µl. One sterilized, unused filter was included as a negative control for each sampling day and sample processing protocol.

### PCR amplification

Metabarcoding was carried out using two mitochondrial primer sets. The vertebrate-specific 12S rRNA primer pair 12SV05 (forward 5′-TTAGATACCCCACTATGC-3′; reverse 5′-TAGAACAGGCTCCTCTAG-3′) amplifies a fragment of approximately 97–103 bp from the 12S rRNA gene (Riaz et al., 2011). Mammals were also targeted using the 16S rRNA primer pair 16Smam1 (forward 5′-CGGTTGGGGTGACCTCGGA-3′) and 16Smam2 (reverse 5′-GCTGTTATCCCTAGGGTAACT-3′), which amplifies a ∼95 bp region of the 16S rRNA gene (Taylor, 1996). Twenty-nine PCR reactions were prepared for each primer pair, including four filter blanks and one PCR negative control prepared with sterile nuclease-free water. Each 20 μL PCR mixture contained 3 μL DNA template, 0.75 U AmpliTaq Gold, 1× Gold PCR Buffer, and 2.5 mM MgCl₂ (all reagents from Applied Biosystems, Thermo Fisher Scientific, Waltham, MA, USA); 0.6 μM each of 5′ nucleotide-tagged forward and reverse primers; 0.2 mM dNTP mix (Invitrogen™, Carlsbad, CA, USA); 0.5 mg/mL bovine serum albumin (BSA; New England BioLabs); and 3 μM human blocker (5′–3′ TACCCCACTATGCTTAGCCCTAAACCTCAACAGTTAAATC–spacerC3 for the 12S rRNA primers (Calvignac-Spencer et al., 2013) and 5′–3′ GCGACCTCGGAGCAGAACCC–spacerC3 for the 16S rRNA primers (Vestheim & Jarman, 2008). For the 12S rRNA primer pair, thermal cycling conditions were: 95 °C for 10 min, followed by 45 cycles of 94 °C for 30 s, 51 °C for 30 s, and 72 °C for 60 s, with a final extension at 72 °C for 7 min. For the 16S rRNA primer pair, cycling conditions were: 95 °C for 10 min, followed by 45 cycles of 95 °C for 12 s, 59 °C for 30 s, and 70 °C for 25 s, with a final extension at 72 °C for 7 min. Amplification was verified using 2% EX pre-cast agarose gels run on the E-Gel™ Power Snap Electrophoresis system (Thermo Fisher Scientific, Waltham, MA, USA). A symmetrical tagged primer system was used for sample multiplexing. Unique 9 bp tags, generated with Barcode Generator (https://github.com/lcomai/barcode_generator), were appended to the 5′ ends of both forward and reverse primers for each primer set. This tag length has been shown to be suitable for multiplexing amplicons in nanopore sequencing (Srivathsan et al., 2024). Tags differed by at least three base pairs from one another.

### DNA Sequencing

The 29 PCR products of each primer set were pooled by taking 15 μl of each product and then cleaned using a 1.5:1 bead-to-DNA ratio with AMPure XP Beads (Beckman Coulter, Germany), followed by resuspension in 50 μL of DNase-free water. Sequencing was performed using the portable temperature-robust MinION Mk1D device. Library preparation was done for each primer pool using the Ligation Sequencing Kit (SQK-LSK114), following the manufacturer’s Ligation Sequencing Kit Amplicons protocol. Both primer pools were sequenced on separate MinION flow cells for approximately 15 hours.

### Database preparation

A reference database was constructed from MIDORI2 (Machida et al. 2023) using the LONGEST nucleotide release (GenBank version 269; MIDORI2_LONGEST_NUC_GB269). The database was filtered to include only Chordata, following best-practice recommendations for vertebrate eDNA studies (Ushio et al. 2023). Quality control steps were applied: sequences with more than ten ambiguous (“N”) nucleotides or exceeding 5% N content were removed. All taxonomic names in the reference database were reconciled against the GBIF Backbone Taxonomy (GBIF.org 2024) through chunked calls (500 records per request), ensuring standardized nomenclature throughout. The curated reference database was reformatted into SINTAX format and indexed using a vsearch unidirectional database (UDB) for rapid similarity searches. Species occurrence within Zambia was verified against a curated vertebrate species list for the country to assess reference database completeness

### Sequence analysis

Raw sequencing reads from all samples and negative controls were basecalled using Dorado v1.0.2 (super-accuracy model: dna_r10.4.1_e8.2_400bps_sup@v4.3.0), retaining reads with a minimum Q score of 10. Basecalled reads were demultiplexed using OBITools4’s obimultiplex command, allowing for a maximum of two errors in the tags (https://git.metabarcoding.org/obitools/obitools4). Reads were then filtered using VSEARCH v2.28.1 to retain sequences of a sequence length between 90–130 bp with a maximum expected error (maxEE) of 0.15 (approximately Q28 for 100 bp reads). Singletons were removed, and globally dereplicated sequences with a minimum abundance of two reads were retained. Chimeric sequences were removed using UCHIME3-denovo in VSEARCH v2.28.1.

We employed Swarm v3.1.0 (Mahé et al. 2015; Mahé et al. 2022) as an Operational Taxonomic Unit (OTU) clustering approach that is based on a local sequence divergence threshold (d). Swarm uses a bottom-up, greedy clustering approach (starting at d=1) that preserves the structure of sequence diversity while avoiding artificial merging of closely related species, which may differ by less than 3% sequence identity in short (∼100 bp) barcode regions. Fastidious mode was enabled to link small clusters to larger clusters in a second-pass analysis, reducing the number of spurious singletons while maintaining biological resolution. Post-clustering curation was performed using mumu (Frøslev et al. 2017, C implementation), which identifies and merges error-derived OTUs based on sequence similarity (vsearch self-matching at 95% identity and 90% query coverage), co-occurrence patterns, and relative abundance ratios. This step reduces inflated diversity caused by sequencing errors and conservative clustering thresholds while preserving genuine sequence variation

### Taxonomic assignment

Initial taxonomic assignment of the OTUs was performed using vsearch --usearch_global against the curated MIDORI2_LONGEST_NUC_GB269 reference database, using relaxed thresholds of ≥90% sequence identity and ≥80% query coverage to retain candidate matches. For each OTU, only hits with the highest percent identity were kept. Among these, hits with bit scores within 99% of the maximum bit score for that OTU were retained, and up to 50 top-scoring matches (including ties) were carried forward for taxonomic annotation. Sequences that did not meet the minimum alignment thresholds were assigned as “unassigned.” For sequences meeting the initial thresholds, a multi-stage assignment framework was applied, where the species list of terrestrial vertebrates known to occur in Zambia is based on checklists retrieved from the SASSCAL portal (2017) (http://data.sasscal.org/metadata/view.php?view=doc_documents&id=3154)

1) OTUs matching known contaminants (12 known domestic/commensal species: *Homo sapiens*, *Bos taurus*, *Canis lupus familiaris*, *Felis catus*, *Sus scrofa*, *Gallus gallus*, *Capra hircus, Ovis aries, Anas platyrhynchos, Meleagris gallopav* and *Equus caballus*) were assigned to the category “contaminant”. Additional “contaminants” were identified from the negative PCR and extraction controls: OTUs detected in at least one negative control but completely absent from all air samples; and OTUs with a read count in a negative control that is larger than the maximum read count in any air sample. 2) If exactly one species from the Zambian vertebrate species list matched at ≥97% identity and ≥90% query coverage, the OTU was assigned to that “unambiguous local” species. 3) If multiple species matched but only one was found within the Zambian species list, the OTU was assigned to that “arbitrated local” species. 4) If exactly one species matched at ≥97% identity and ≥90% coverage, but was not on the Zambian list, the OTU was assigned to “unambiguous nonlocal” species and was removed for downstream analysis. 5) If the best match was a non-local species, but a species of the same genus was present on the Zambian species list and absent from the MIDORI2 database, the OTU was flagged as “nonlocal_local_congener_missing” and kept at the genus level, highlighting potential gaps in the reference database. 6) If multiple best hits resolved only to genus level, the OTU was retained at genus level (“ambiguous_genus”). 7) OTUs not meeting any of the above criteria remained unassigned. This assignment framework allows the Zambian species list to serve as a guide for intelligent taxonomic arbitration and quality control, rather than a hard filter that discards potentially valuable detections.

### Species-accumulation curves

Species richness accumulation curves were generated with iNEXT rarefaction and extrapolation (Hsieh et al. 2016), using incidence frequency data (q=0 for species richness) with a target coverage of 98%. Separate curves were generated to illustrate richness accumulation across sampling days (pooling across all samplers per day) and across samplers (pooling across all days per sampler).

## Results

All air sampling metadata is available in Table S1. Subsequent DNA concentration and purity measurements can be found in Table S2 (Materials and Methods). For the 12S rRNA marker, 21.3M raw reads were generated, of which 94.8% (20.2M) were successfully demultiplexed, with 14,573–4,711,550 reads per sample. For the 16S rRNA marker, 11.64M raw reads were obtained, of which 25.0% (2.91M) were demultiplexed, with 7,187–296,959 reads per sample (Table S3). After quality filtering and chimera removal, 34,330 (12S) and 3,254 (16S) unique sequences were retained for sequence clustering. Swarm v3.1.0 denoising (d=1, fastidious mode) identified 3,215 (12S) and 319 (16S) OTUs. Following mumu post-clustering curation, 989 (12S) and 195 (16S) OTUs were retained (Materials and Methods).

Following our multi-stage taxonomic assignment framework (Materials and Methods), OTUs were categorized into several taxonomic groups (Table S4). In the 12S rRNA dataset, 98 OTUs were classified as “unambiguous_local”, 57 as “ambiguous_genus”, 30 as “unambiguous_nonlocal”, 20 as “nonlocal_local_congener_missing”, and 2 as “unassigned”, resulting in a total of 207 assigned OTUs. In the 16S rRNA dataset, 45 OTUs were assigned to “unambiguous_local”, 8 to “ambiguous_genus”, and 12 to “unambiguous_nonlocal”, resulting in a total of 65 assigned OTUs. No OTUs were categorized as “arbitrated_local” in either dataset. The OTUs classified as “unambiguous_nonlocal” were excluded from all downstream analyses.

In the negative controls (n = 5; four filter blanks and one PCR negative control), 17 (1) of the OTUs of the 12S (16S) rRNA dataset had at least as many sequencing reads in a negative control as they had in any of the air samples, and were therefore removed from downstream analysis. No OTU only occurred in only the negative controls (Materials and Methods). One out from the 16S dataset was removed under the same criterion. No OTUs were found exclusively in negative controlsRemoval of contaminants resulted in 71 (12S) and 39 (16S) OTUs assigned to “unambiguous_local” species, nine (12S) OTUs classified as “ambiguous genus” were included at the genus level and additional 18 OTUs (12S) were flagged as “nonlocal_local_congener_missing” and kept at the genus level. These were cases where a Zambian congener is present on the species list but lacked a representative sequence in MIDORI2 GB269 pointing to reference database gaps and are candidates for future database curation.

These remaining OTUs were assigned to 120 Zambian terrestrial vertebrate taxa (12S: 96; 16S: 40). Within the 12S rRNA dataset, 59 bird (Aves), 31 mammal (Mammalia), three reptile (Reptilia), and three amphibian (Amphibia) taxa were detected. Within the 16S rRNA dataset, 37 mammal and three amphibian taxa were detected. Sixteen mammal taxa were detected by both markers. Taxonomic detections were unevenly distributed across orders (Figure 2A). Among birds, Passeriformes was the most taxonomically diverse order, with 21 taxa detected and a total of 243 detections. Other taxa-rich bird orders included Pelecaniformes (7 taxa; 57 detections) and Galliformes (5 taxa; 89 detections). Among mammals, Artiodactyla was the most species-rich order, comprising 15 taxa and 126 total detections. This was followed by Rodentia (12 taxa; 71 detections) and Carnivora (10 taxa; 52 detections). At the class level, birds exhibited the highest taxonomic richness.

**Figure 2.**
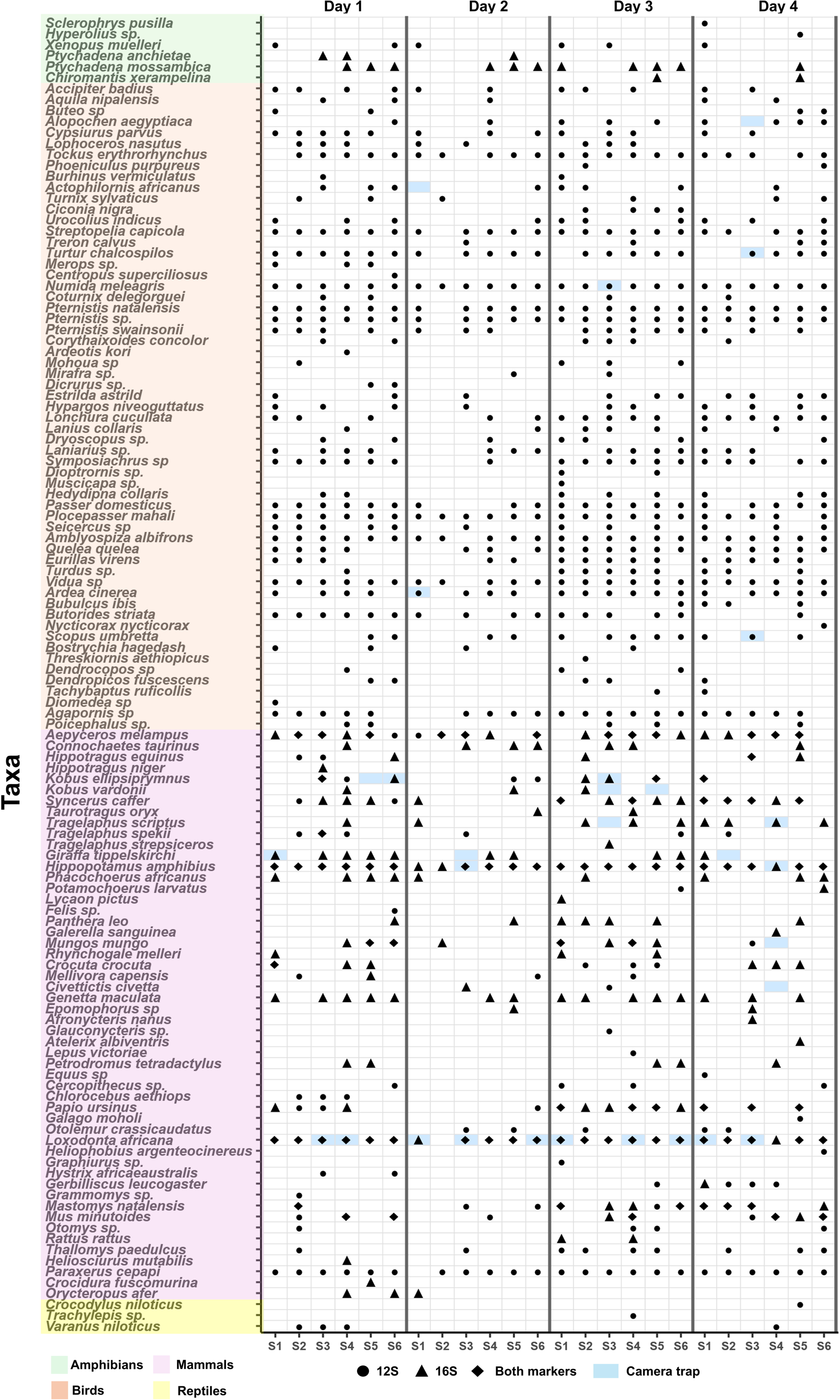
(A) Number of taxa detected sorted by terrestrial vertebrate order. (B–C) Species-accumulation curves for airborne eDNA and camera traps generated using incidence-frequency iNEXT analyses. Panel (B) shows day-level accumulation (pooling six locations per day; n = 4 days), and panel (C) shows sampler-level accumulation (pooling four days per location; n = 6 locations). Solid lines represent rarefaction and dashed lines extrapolation; shaded areas indicate 95%-confidence intervals (CI). Right-side labels show richness standardized to 98% sample coverage (SC = 0.98). (D) Detection-frequency distribution of terrestrial vertebrate taxa across classes; taxa detected by both markers are counted twice

Taxa detections were strongly right-skewed, with most taxa detected infrequently (Figure 2D; Figure S9). Of the 120 taxa detected, 26 taxa (21.7%) were recorded only once, 37 taxa (30.8%) were recorded 2–5 times, 25 taxa (20.8%) were recorded 6–10 times, and 20 taxa (16.7%) were recorded 11–20 times. Only 12 taxa (10.0%) were recorded more than 20 times, including 2 taxa detected more than 40 times. At the class level, birds exhibited the highest mean number of detections per taxon (mean = 9.9, SD = 7.6). Mammals averaged 5.8 detections per taxon (SD = 6.2) in the 12S rRNA dataset and 6.0 detections per taxon (SD = 6.3) in the 16S rRNA dataset. Amphibians and reptiles showed lower overall detection frequencies. Amphibians averaged 2.7 detections per taxon (SD = 2.9) for 12S rRNA and 5.3 detections per taxon (SD = 4.9) for 16S rRNA. Reptiles had a mean of 2.0 detections per taxon (SD = 1.7).

The most frequently detected taxa overall were hippopotamus (*Hippopotamus amphibius*; 45 detections) and African elephant (*Loxodonta africana*; 44 detections), followed by impala (*Aepyceros melampus*; 21 detections). Among mammals, the bush squirrel (*Paraxerus cepapi*; 23 detections) was also frequently detected. Among birds, the helmeted guineafowl (*Numida meleagris*; 24 detections) was the most frequently detected species, followed by several passerines, including *Amblyospiza albifrons* (23 detections) and *Plocepasser mahali* (23 detections) (Figure 3).

**Figure 3.**
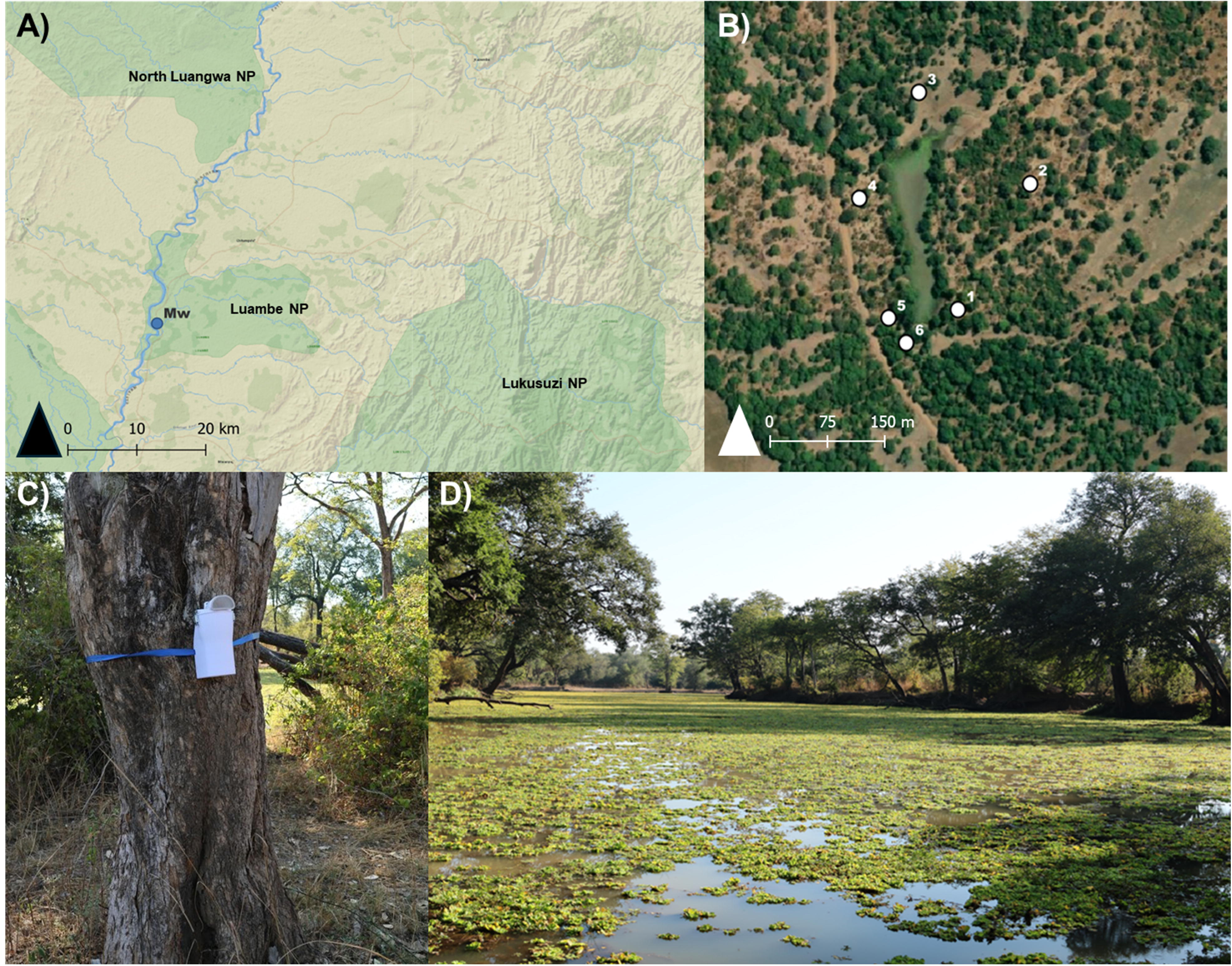
Terrestrial vertebrate taxa detections across four sampling days (panels) and six samplers per day (columns). Airborne eDNA detections are indicated by circles (12S rRNA), triangles (16S rRNA), and diamonds (both markers), while light blue tiles represent camera trap-based detections.

Species-accumulation curves indicated that the combined sampling effort (24 biological samples across four sampling days and six sampler locations) approached high sample completeness but did not fully saturate species richness (Figures 2B-C). At the day level (pooling all six samplers per day; n = 4 days), the observed richness was 120 taxa. When standardized to 98% sample coverage (SC = 0.98), estimated richness increased to 136.6 taxa (95%-confidence interval (CI): 120.6–152.5). Similarly, at the sampler level (pooling all four days per sampler; n = 6 samplers), richness at SC = 0.98 was estimated at 140.7 taxa (95%-CI: 115.4–166.0).

Day-level species-accumulation (Figure 2B, Figure S7) showed a rapid initial increase in detected richness followed by smaller but consistent gains with additional sampling days. When all samplers were combined, 87 taxa were detected on Day 1, representing 72.5% of the total 120 taxa. Richness increased to 93 taxa (77.5%) on Day 2, and 110 taxa (91.7%) on Day 3. These results indicate that a single day of sampling captured most detectable taxa, while subsequent days contributed additional but progressively fewer new detections.

Sampler-level species-accumulation (Figure 2C; Figure S8) revealed a similar pattern. When pooling all days, a single sampler detected 74 taxa (61.7% of the total 120 taxa). Adding a second sampler increased richness to 89 taxa (74.2%), three samplers to 101 taxa (84.2%), four samplers to 108 taxa (90.0%), and five samplers to 115 taxa (95.8%). Thus, although individual samplers captured a substantial proportion of the total richness, each additional sampler consistently contributed new detections, with the largest gains occurring when increasing from one to two and three samplers.

Camera traps recorded 17 species, all of which were also detected using airborne eDNA except for one species, the lilac-breasted roller (*Coracias caudatus*) (Table S6). One mongoose (Herpestidae) detected by camera trapping could not be identified to species level whereas airborne eDNA resolved three mongoose species: the banded mongoose (*Mungos mungo*), Meller’s mongoose (*Rhynchogale melleri*), and the slender mongoose (*Galerella sanguinea*). At the finer scale of species-by-day detections, camera traps recorded 30 events, of which 20 were matched by airborne eDNA at the same location on the same day (Figure 3).

Rarefaction–extrapolation analyses of the camera-trap detections similarly indicated that sampling did not reach high completeness within the study period (Figures 2B-C). At the day level (pooling all six cameras per day; n = 4 days), observed richness of 17 taxa corresponded to an estimated richness of 27.7 taxa at 98% sample coverage (95% CI: 0.0–60.4). At the sampler level (pooling all days per camera; n = 6 cameras), richness at SC = 0.98 was estimated at 30.0 taxa (95% CI: 8.2–51.8).

## Discussion

We demonstrate that an on-site airborne eDNA approach can capture a broad and taxonomically diverse community of vertebrates in a subtropical savanna ecosystem over a short, four-day sampling period. Airborne eDNA has been proposed as a scalable tool for biodiversity monitoring, both temporally and spatially, with the potential to survey large geographic areas and track community dynamics over extended periods (Tulloch et al., 2025; Sullivan et al., 2025; Tournayre et al., 2025). Our results show that, in addition to long-term scalability, airborne eDNA can also provide a rapid and informative snapshot of biodiversity within a short sampling window. By integrating the portable and affordable MinION sequencing device into our air eDNA framework, we further provide means of generating and interpreting eDNA data rapidly on-site, obviating the need for expensive and time-consuming sample storage and transport, and potentially increasing accessibility in LMICs where many regions of high conservation importance are located (Pomerantz et al., 2018; Urban et al., 2023; Gygax et al., 2025).

In our study in air eDNA study in the Luangwa Valley in Zambia, we detected 120 vertebrate taxa that could be assigned to local species/genera across the four major terrestrial vertebrate classes (mammals, birds, reptiles and amphibians). The detected taxa spanned multiple vertebrate classes and encompassed broad ecological diversity, demonstrating that airborne eDNA can recover taxonomically and ecologically diverse vertebrate assemblage, supporting its use as an additional tool for community-level biodiversity assessments in different terrestrial ecosystems such as savannas, temperate and tropical forests (e.g. Lynggaard et al., Searle, 2022; Craine et al., 2025).

In our study, birds and mammals showed the highest representation, which may reflect their higher local biomass, greater movement across the landscape, and therefore higher relative DNA shedding rate, all of which have been suggested to increase vertebrate detection probabilities in airborne eDNA surveys. (Lynggaard et al., 2022; Nordstrom et al., 2022; Newton et al., 2025; Sullivan et al., 2025; Tulloch et al., 2025). The comparatively low detection rates of amphibians and reptiles could therefore result from low shedding rates, low biomass and mobility, and their partially semi-aquatic lifestyle. However, we cannot rule out technical effects of primer amplification bias and lack of PCR replicates on the detection rate of the different vertebrate taxonomic groups (Wang et al., 2023; Craine et al., 2025).

The vertebrate orders with the highest taxonomic richness in our dataset were Artiodactyla (even-toed ungulates) and Rodentia (rodents) among mammals, and Passeriformes (passerine birds) among birds. The strong representation of ungulates in our airborne eDNA data, including the frequent detection of hippopotamus (*H. amphibius*), elephants (*L. africana*), and impala (*A. melampus*), is consistent with the known high diversity and biomass of medium- to large-bodied herbivores in the Luangwa Valley, with more than 30 species reported in Zambia (Ndhlovu & Balakrishnan, 1991; Anderson et al., 2016). The frequent detection of smaller rodent and bird taxa might be consistent with allometric scaling relationships described by Yates et al. (2021), who argued that smaller organisms shed disproportionately higher amounts of eDNA per unit biomass than larger organisms. For example, studies suggest that rodents in African savannas may reach densities where their collective energy consumption rivals that of large ungulates (Keesing, 2000), contributing substantially to ecosystem biomass and energy turnover despite their small body size.

The strong representation of rodents in our airborne eDNA dataset is also consistent with their known local diversity in Zambia (more than 70 species) and their high abundance in Luambe National Park specifically—according to past line-transect surveys (Anderson et al., 2016). These surveys especially found a high abundance of the bush squirrel (*P. cepapi*), which accounted for 249 of 315 recorded rodent observations; *P. cepapi* was the third most frequently detected mammal in our airborne eDNA data. Because rodents are small, fast-moving, often cryptic, and generally not detectable by acoustic methods, they may be overlooked by visual surveys and conventional camera trapping, while live trapping with devices such as Sherman traps is labor-intensive and invasive (Šklíba et al., 2019; Thomas et al., 2020; Hopkins et al., 2024). While conventional monitoring methods remain valuable because they can provide information on activity and abundance, our results suggest that airborne eDNA can offer a complementary, non-invasive approach for efficient monitoring rodents.

The strong representation of passerine birds in our airborne eDNA dataset is consistent with Passeriformes being the largest avian order, comprising more than 300 species in Zambia and dominating many terrestrial bird communities in abundance, diversity, and metabolic activity (Callaghan et al., 2021). Among the most frequently detected bird species was the non-passerine guineafowl (*N. meleagris*), whose high detection frequency likely reflects its gregarious behavior and high local abundance in the area (pers. obs.).

Camera traps provided an independent means of validating taxon detections, and here resulted in 17 vertebrate taxon detections. Our matched airborne eDNA data detected nearly all these taxa (16 out of 17), supporting its use as a sensitive and complementary tool for terrestrial vertebrate monitoring. Camera trap detections are influenced by factors such as camera placement, orientation, and species-specific movement behavior, which can influence detection probabilities and may result in some taxa being recorded more frequently than others (Kolowski & Forrester, 2017; DeWitt & Cocksedge, 2023; Kalan et al., 2023). While camera traps provide direct, localized evidence of species presence, broader representation of the vertebrate community generally requires increased spatial replication and longer deployment periods (Si et al., 2014; Searle et al., 2022). In contrast, airborne eDNA sampling may integrate biological signals across space and time, potentially complementing camera-based community level monitoring (Warmer et al., 2025).

Species-accumulation curves showed that the combined sampling effort (24 biological samples across four sampling days and six sampler locations) approached high biodiversity completeness, with observed richness of 120 taxa at an estimated richness of ∼136 taxa at 98% sample coverage. Although additional sampling efforts would likely detect further taxa, gains are expected to be moderate relative to the richness already captured. Importantly, most detectable diversity was captured early in the sampling period: Temporally, 71.4% of the taxa were detected on the first day alone, with subsequent days contributing progressively fewer new taxa. By the third day, 92% of the total richness had been recorded. Spatially, a single sampler detected nearly 59% of all taxa, and the largest increase in richness occurred when moving from one to two or three samplers, after which gains diminished. These patterns indicate that airborne eDNA can efficiently capture a substantial fraction of local vertebrate diversity with limited temporal effort, while additional sampling efforts might primarily enhance the detection of less common or spatially heterogeneous taxa. Together, these results suggest that monitoring strategies could prioritize broader spatial coverage with shorter deployment durations when the goal is to obtain representative biodiversity snapshots. At the same time, detection of rarer or less abundant species may benefit from methodological optimization, such as the use of multiple genetic markers and increased PCR replication per sample (Craine et al., 2025) to enhance sensitivity without substantially increasing sampling efforts.

In a water eDNA study conducted at the same waterhole at a similar time, we evaluated water-based eDNA alongside camera trap detections (Gygax et al., 2025). Briefly, water eDNA data detected 10 vertebrate taxa, including only 4 of the 7 taxa recorded by camera traps. In our airborne eDNA study, on the other hand, airborne DNA detected 120 vertebrate taxa, including 16 of the 17 taxa recorded by camera traps. While differences in study design and sampling efforts prevent a direct quantitative comparison between these water and air eDNA studies, this might suggest airborne eDNA recovering a broader spectrum of terrestrial vertebrates than water eDNA. Airborne eDNA can further capture signals over broader spatial scales, is more scalable, and less dependent on animal ecology and behavior (Tulloch et al., 2025). In contrast, water eDNA might provide a more localized signal, and its ecological dynamics are currently better understood (Harrison et al., 2019; Kirtane et al, 2023; Schenekar et al., 2024). The most appropriate biodiversity monitoring strategy therefore depends on the ecological question and target taxa, and integrating airborne eDNA, water eDNA, and camera trapping can provide a more complete assessment by combining multiple independent data sources (Broadhurst et al., 2025, Warmer et al., 2025).

Airborne eDNA—similar to other eDNA approaches—is subject to errors arising at different stages of the workflow. Tulloch et al. (2025) proposed a framework distinguishing four major error types across two stages: detection (during sample collection) and identification (during data analysis). These include false-negative detections, where DNA is present in the environment but not captured; false-positive detections, where DNA is correctly identified but originates from outside the target area; false-negative identifications, where DNA is captured but cannot be accurately identified; and false-positive identifications, where DNA is misidentified as the wrong species.

False-negative detections are difficult to confirm. In our study, there were several instances where a species detected by camera trapping on a given day was not detected by the corresponding air sampler (although the reverse pattern was more common). It is further likely that some taxa were missed by both approaches. In addition, DNA may have been successfully captured but failed to amplify, as frequently observed across PCR replicates (Craine et al., 2025). False-positive detections are particularly challenging in airborne DNA studies because of the potential for long-range DNA transport and contamination (Tulloch et al., 2025). DNA might therefore be correctly identified but originate from outside the focal study area, complicating ecological interpretation; a distinction between long-distance transport and local presence might remain inherently difficult in airborne eDNA data interpretation.

False-negative identifications can here be inferred from the—however limited—overlap between genetic markers: We detected 31 mammal taxa with the 12S rRNA marker and 37 with the 16S rRNA marker, with only 16 taxa shared between markers. This suggests that some taxa were present in the samples but were detected or resolved by only one marker, likely due to primer bias or amplification stochasticity. Incomplete reference databases aggravate this issue. Here, reference coverage for Zambian terrestrial vertebrates was incomplete for both markers: in the Zambia species list, 44.4% of species and 79.2% of genera were represented in the 12S rRNA reference database, and 46.0% of species and 78.1% of genera in the 16S rRNA database. Such incomplete reference coverage increases the risk of both false-negative and false-positive identifications. In our dataset, 20 OTUs matched a locally present genus for which no local species-level reference sequence was available. Additionally, 20 OTUs could not be resolved beyond the genus level. While the incorporation of local species lists can help reduce misidentifications, care has to be taken when designing such lists and their spatial resolution.

Using a Zambia-wide species list, we, for example, classified three primate taxa (*Papio ursinus*, *Chlorocebus aethiops*, and the genus *Cercopithecus* sp.) as unambiguous local taxa detections. However, the Luangwa Valley only hosts populations of *Papio cynocephalus* and *Chlorocebus pygerythrus*, which suggests that our genetic markers did not have sufficient resolution to reliably distinguish between closely related species, while long-distance DNA transport can also not be ruled out. *Cercopithecus* sp. Is further not known to occur in the Luangwa Valley, but long-distance DNA transport can again not be ruled out.

Bioinformatic decisions strongly impact these major error types. Applying stringent filtering or merging thresholds (e.g., removing OTUs with fewer than 100 reads or those occurring in few samples) can reduce false-positive identification, but can also remove rare but true detections and increase the number of false-negative identifications. For example, we detected Nile crocodile (*Crocodylus niloticus*), a species known to occur in the area, in only one sample and at low read counts, which would have been excluded if stricter filtering parameters had been applied. Similarly, increasing the sequence identity threshold for taxonomic assignments to more than 97% would have excluded the detection of *P. cepapi*, a locally abundant species whose intraspecific genetic diversity might not be sufficiently represented in the reference database. We therefore conclude that no single parameter set is optimal across all taxa, ecosystems, and markers, and careful interpretation of any taxa detection is still required when it comes to eDNA data analysis. Multi-stage classification and arbitration frameworks such as the one applied in this study integrate taxonomic assignment ambiguity, local species knowledge, negative control screening, and reference availability, and can therefore help improve detection and identification errors. Our study demonstrates that on-site airborne eDNA can recover a taxonomically and ecologically diverse vertebrate community within a short sampling period in a subtropical savanna ecosystem. Although methodological limitations remain, especially regarding spatial resolution, reference database completeness, and accurate taxonomic assignments, our results highlight the practical utility of airborne eDNA as a scalable, efficient, and rapid biodiversity monitoring tool.

## Conclusion

We demonstrate that on-site airborne eDNA can accurately detect the presence of taxonomically and ecologically diverse terrestrial vertebrates in a subtropical savanna ecosystem. We captured taxa across all four terrestrial vertebrate classes and showed substantial concordance with camera trapping-based detections. With further methodological advances, airborne eDNA is likely to become a valuable component of integrated biodiversity monitoring frameworks, including in remote and conservation-priority regions such as the Luangwa Valley.

## Data Accessibility Statement

Sequencing data from this study have been submitted to NCBI under BioProject PRJNA1435361. The associated BioSample accessions are SAMN56444883-SAMN56444911. SRA run accession numbers are currently pending and will be updated in a later version of the manuscript. Analysis code is available at https://github.com/dmgr90/Zambia_air/tree/main.

## Acknowledgements

We thank the Department of National Parks and Wildlife for this fruitful collaboration. This 434 research was carried out with their authorization under the research permit NPW/8/27/1.

## Notes

### Competing Interest Statement

The authors have declared no competing interest.

