## Supplementary figures and images for "Monitoring terrestrial vertebrates with airborne DNA in the Luangwa Valley, Zambia"

### Supplementary Figure S7

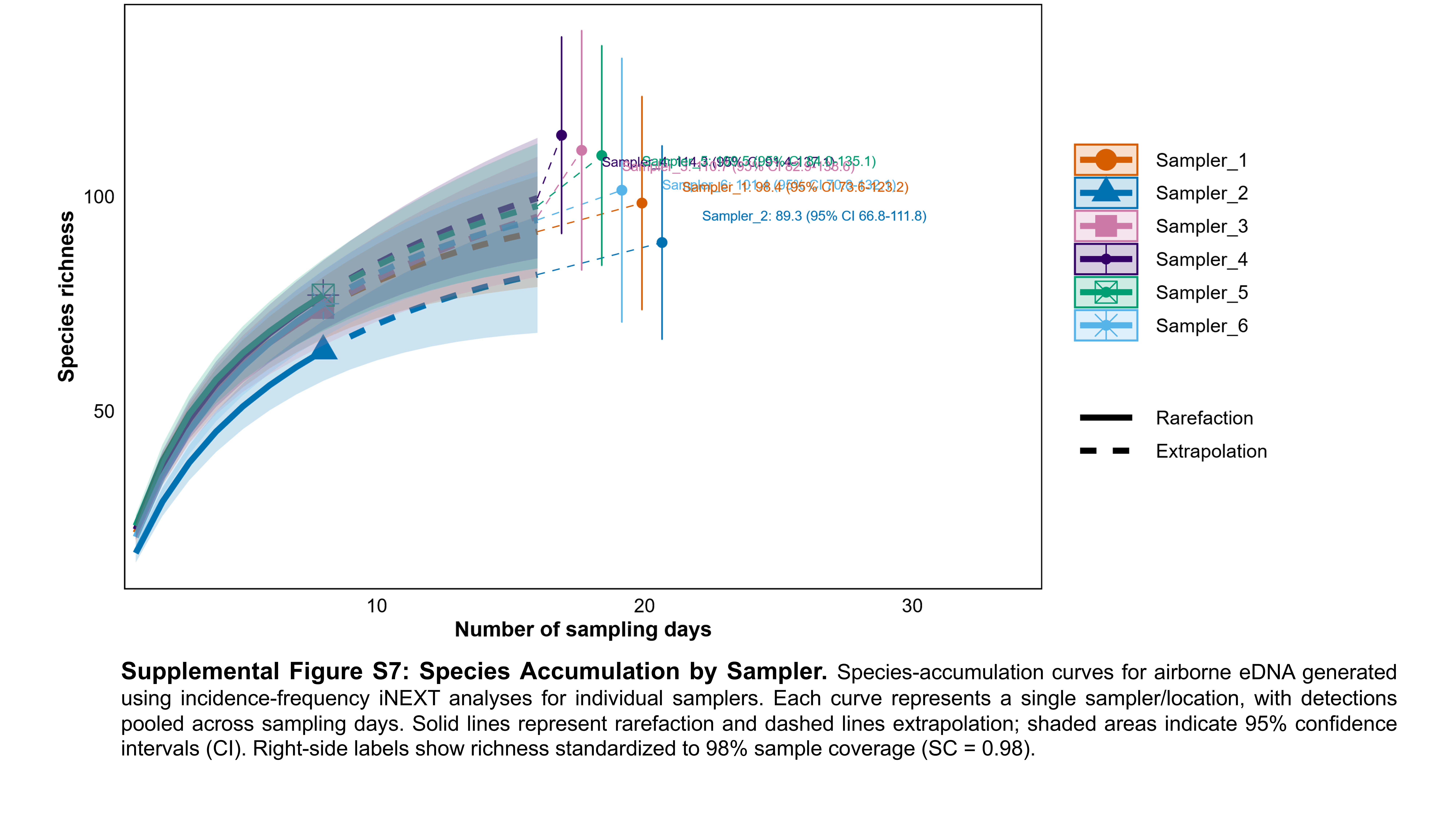

### Supplementary Figure S8

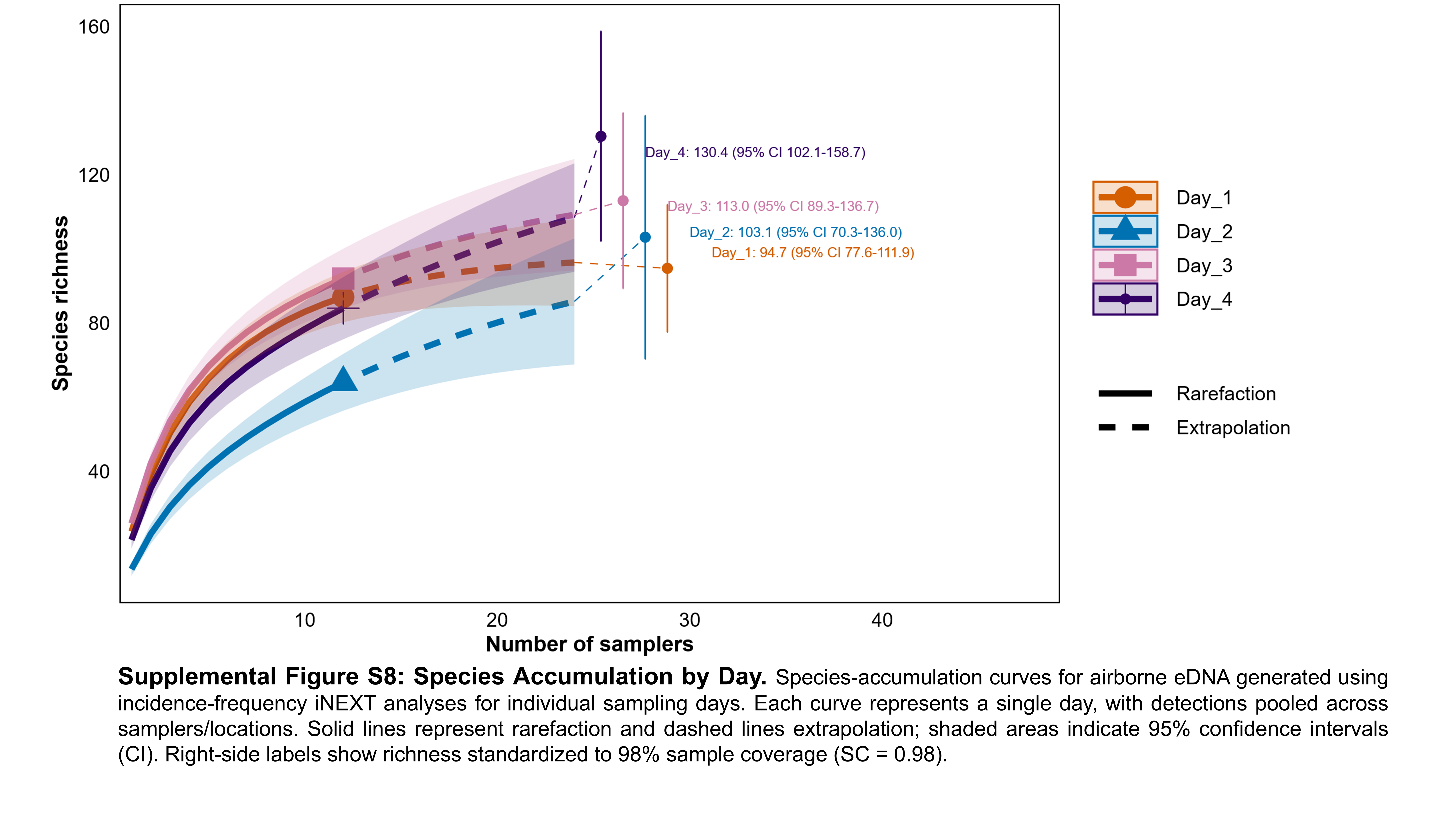

### Supplementary Figure S8

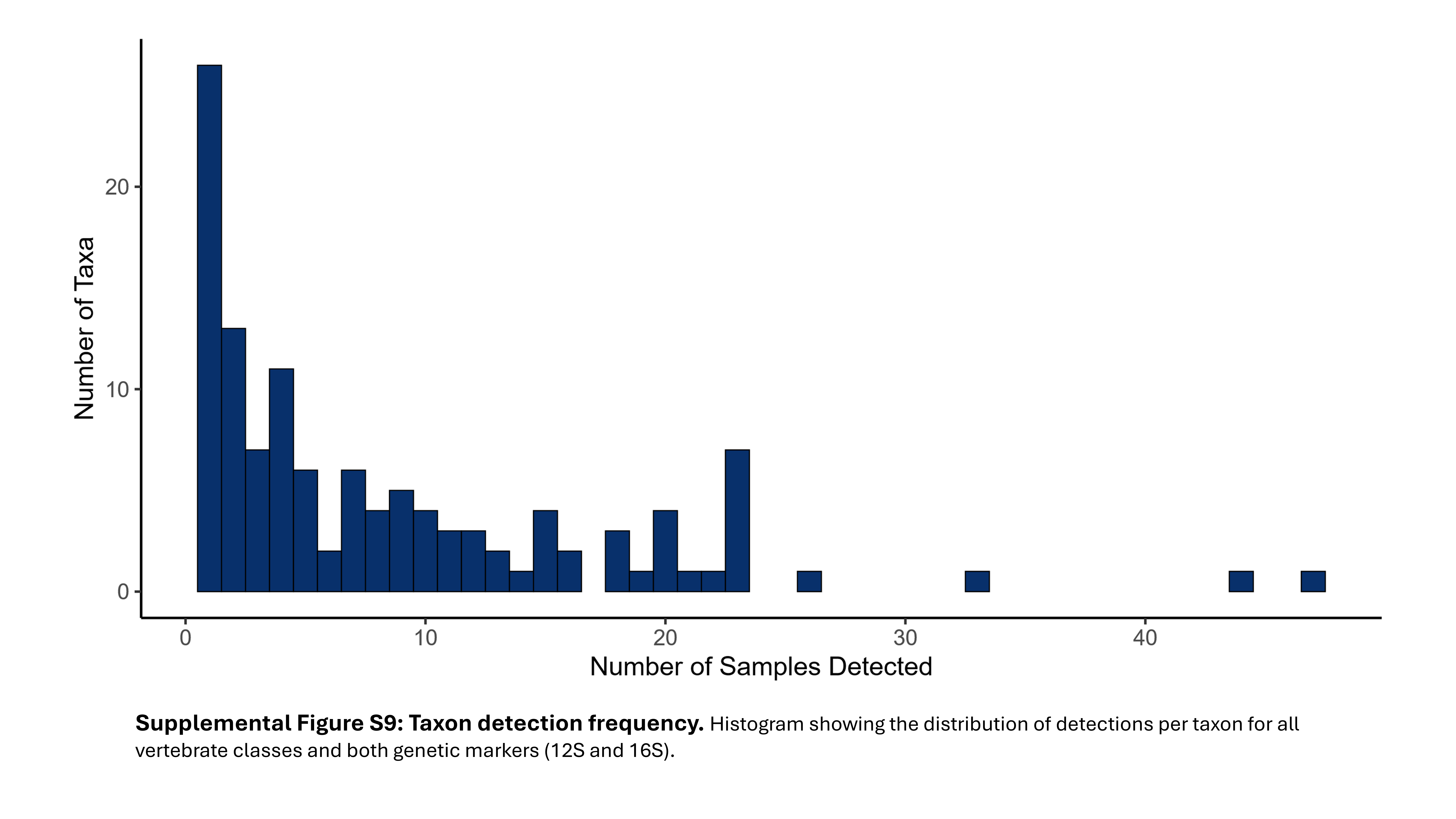
